# Learning and transfer of working memory gating policies

**DOI:** 10.1101/144543

**Authors:** Apoorva Bhandari, David Badre

**Affiliations:** Cognitive, Linguistics, & Psychological Science, Brown University, Providence, USA; Brown Institute for Brain Sciences, Brown University, Providence, USA

## Abstract

Abstract knowledge about the tasks we encounter enables us to rapidly and flexibly adapt to novel task contexts. Previous research has focused primarily on abstract rules that leverage shared structure in stimulus-response (S-R) mappings as the basis of such task knowledge. Here we provide evidence that working memory (WM) gating policies – a type of control policy required for internal control of WM during a task – constitute a form of abstract task knowledge that can be transferred across contexts. In two experiments, we report specific evidence for the transfer of selective WM gating policies across changes of task context. We show that this transfer is not tied to shared structure in S-R mappings, but instead in the dynamic structure of the task. Collectively, our results highlight the importance of WM gating policies in particular, and control policies in general, as a key component of the task knowledge that supports flexible behavior and task generalization.

## Introduction

Humans display remarkable cognitive flexibility in novel task environments (McClelland, 2009). Given only verbal instruction, we rapidly adapt to new tasks, often achieving asymptotic levels of performance within just a few trials (Ackerman, 1988; Bhandari & Duncan, 2014; Ruge & Wolfensteller, 2010; Wolfensteller & Ruge, 2011). Such rapid adaptation relies, in part, on abstract task knowledge transferred from prior experience with other tasks. Abstract task knowledge captures regularities in the space of task environments, and can thus speed up learning in the new environment by reducing the size of the learning problem (Botvinick, Niv, & Barto, 2009; Cole, Etzel, Zacks, Schneider, & Braver, 2011; Collins & Frank, 2013; Gershman & Niv, 2010).

What form does such abstract task knowledge take? The vast majority of prior studies seeking to address this question have focused on rules, task-sets, or stimulus-response (S-R) mappings as the basis of task knowledge. In these frameworks, abstract rules generalize prior knowledge and thus constrain the (usually) very large space of stimulus-response-outcome contingencies afforded by a novel task environment (Badre, Kayser, & D'Esposito, 2010). Such rules can both be instructed (Cohen-Kdoshay & Meiran, 2007, 2009; Cole, Bagic, Kass, & Schneider, 2010; Meiran, Pereg, Kessler, Cole, & Braver, 2015; Ruge & Wolfensteller, 2010) or transferred from prior experiences (Cole et al., 2011; Collins & Frank, 2013) to rapidly enable successful behavior in novel environments.

The implementation of a task, however, requires more than just the knowledge of stimulus-response-outcome contingencies. Even the simplest everyday task environments have dynamical structure, with events unfolding in a specific order, and with specific timing (Radvansky & Zacks, 2014). To achieve task goals in a dynamic task environment, then, one must also learn an *internal control policy* or *task model* aligned to the task’s dynamic structure for the moment-by-moment control of internal cognitive processing (Bhandari & Duncan, 2014; Duncan et al., 2008). Such implementational control policies are not typically communicated via instruction and must be discovered and implemented “on the fly”, through task experience.

In this paper, we ask whether control policies are themselves a form of abstract task knowledge that, like rules, can be transferred to novel task contexts. Just like different real-world tasks often share stimulus-response-outcome contingencies, they also share other forms dynamic structure (Botvinick, Weinstein, Solway, & Barto, 2015; Schank & Abelson, 1977). Such shared structure affords an opportunity for generalization of internal control policies. Instead of learning new control policies from scratch, humans may build repertoires of internal control policies that are re-used in novel tasks.

We operationalize this question within the domain of working memory (WM) control – i.e. the selective use of working memory. WM control has been extensively analyzed within the *gating framework,* in which access to WM is controlled by a set of input and output gates (Chatham & Badre, 2015; O'Reilly & Frank, 2006; Todd, Niv, & Cohen, 2009). The contents of WM can be selectively updated by operating an input gate that determines whether stimulus information can enter WM. Similarly, operating a selective output gate allows WM to selectively influence downstream. Learning to perform a WM task, therefore, involves learning a *gating policy* for operating input and output gates in a moment-by-moment, task-appropriate manner (Frank & Badre, 2012). In the context of WM, a gating policy is an example of a control policy that must be aligned to the dynamic structure of the task. By learning such WM gating policies and transferring them across task contexts, humans may be able to exploit regularities in the dynamic structure of tasks.

To test this possibility, we adopt the 2^nd^ order WM control task employed by Chatham, Frank, and Badre (2014). In their task, participants saw a sequence of three items on every trial, one of which specified a context. The context signaled which of the other two items in the sequence was the target item. Critically, there were two kinds of task structures – ‘context first’ (CF) trials, on which the first item in the sequence was the context item, and ‘context last’ (CL) trials, in which the last item in the sequence was the context item. CF and CL trials afford the use of different WM gating policies. On a CL trial, subjects had to employ a ‘selective output-gating policy’ that allowed the storage of both lower level items in WM (a non-selective input-gating operation), and the retrieval of the target item for guiding response selection (a selective output-gating operation). On a CF trial, while a similar selective output-gating policy could be employed, a more efficient ‘selective input-gating policy’ was possible. Such a policy would enable proactive coding of the contextual cue in WM, followed by selective input-gating of only the relevant lower-level item contingent on context. This allows a reduction in both, WM load, and interference from the competing non-target during response selection. Indeed, Chatham et al. (2014) presented evidence that CF and CL trials are treated differently and that well-trained subjects employ selective input-gating policies on CF trials to improve performance relative to CL trials on which the selective output-gating policy is required.

In the context of this WM control task, we ask whether selective gating policies learned in one task setting are transferred to a novel task setting. For instance, subjects exposed to an environment with only CL trials would learn a selective output-gating policy. Would this policy transfer to a new block with CF trials? In Experiment 1 we find a pattern of transfer effects that support the hypothesis that a previously learned gating policy influences initial behavior in a novel setting. We replicate these findings in Experiment 2. In addition, we provide evidence that transferred gating policies are dissociable from S-R mappings and have a much larger influence on subsequent behavior. We interpret these findings as evidence that internal control policies comprise an important form of structural task knowledge that supports behavior in novel situations.

## Experiment 1

### Methods

#### Participants

85 adult, right-handed participants (34 males, 51 females; age-range: 18-30, M = 21.4, SD = 2.7) from the Providence, RI area were recruited to take part in a computer-based behavioral experiment. We endeavored to collect between 18-20 participants in each of four groups based on approximate effect sizes suggested by pilot data. 1 participant was excluded for prior neurological injuries, 3 were excluded as they were on psychoactive medication. 5 participants were excluded because of low performance (< 70% accuracy) on the task. This left 76 participants (30 males, 46 females; age-range: 18-29, M = 21.3, SD = 2.6). Participants were randomly assigned to four groups of 19 each and there were no group differences in age or gender ratio (*p* > 0.250). We subsequently recruited another 38 participants to serve as additional control groups (17 males, 21 females; age-range: 18-30, M = 22, SD = 3.9). All participants had normal or corrected-to-normal vision, and no reported neurological or psychological disorders. All participants gave informed, written consent as approved by the Human Research Protections Office of Brown University, and they were compensated for their participation.

#### Apparatus

Experiments were conducted on a computer running the Mac OSX operating system. The stimulus delivery program was written in MATLAB 2013b, using the Psychophysics Toolbox. Responses were collected via a standard keyboard. All analyses were carried out in MATLAB 2013b and SPSS 22.

#### Task and Experiment Design

Participants were instructed to perform a 2^nd^ order working memory control task (Figure 2). On each trial, they saw a sequence of three items on the computer screen: a number (11 or 53), a letter (A or G), and a symbol (π or ⊙). The number served as a higher-level contextual cue, which specified the lower level items (letter or symbol) that would be the target on each trial. The relationships between the contextual cues and the lower level items were specified by the rule trees shown in Figure 2(a). Participants were asked to memorize these rule trees before the task began, and were given an opportunity to review the trees for as long as they wanted at the beginning of each block.

**Figure 1:**
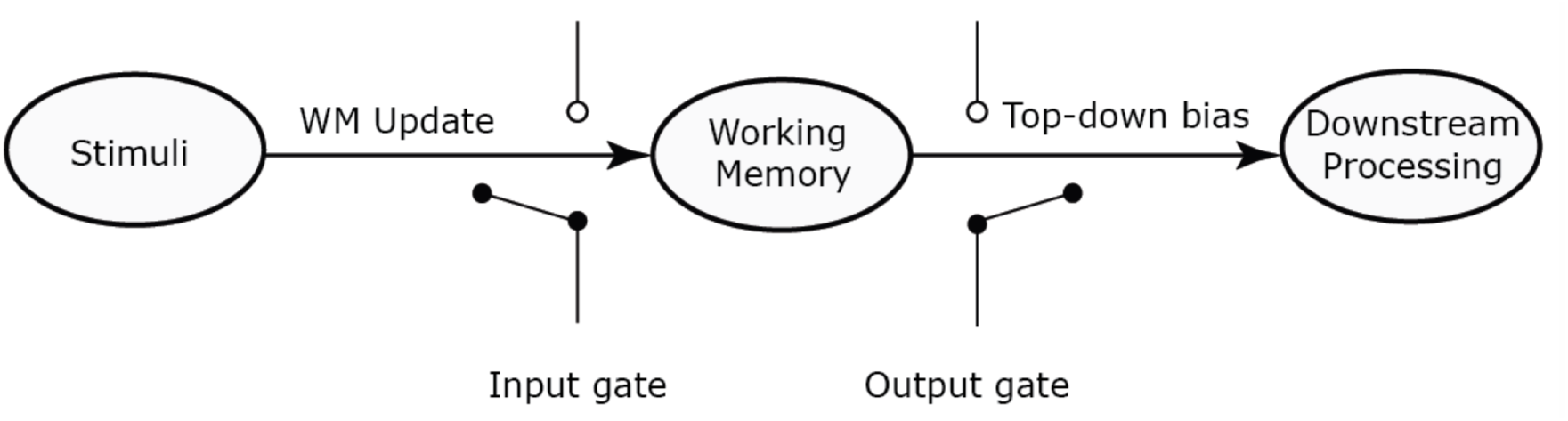
Simple model of working memory control within the gating framework. Access to WM is controlled via the operation of an input gate that determines whether a stimulus is updated into WM. On the other side, an output gate controls whether or not information within WM can influence on behavior. Two broad classes of policies can be distinguished for selective use of WM – ones that achieve selection via selective input-gating, and ones that rely on selection via output-gating. Adapted from Hazy, Frank, and O'Reilly R (2007).

**Figure 2:**
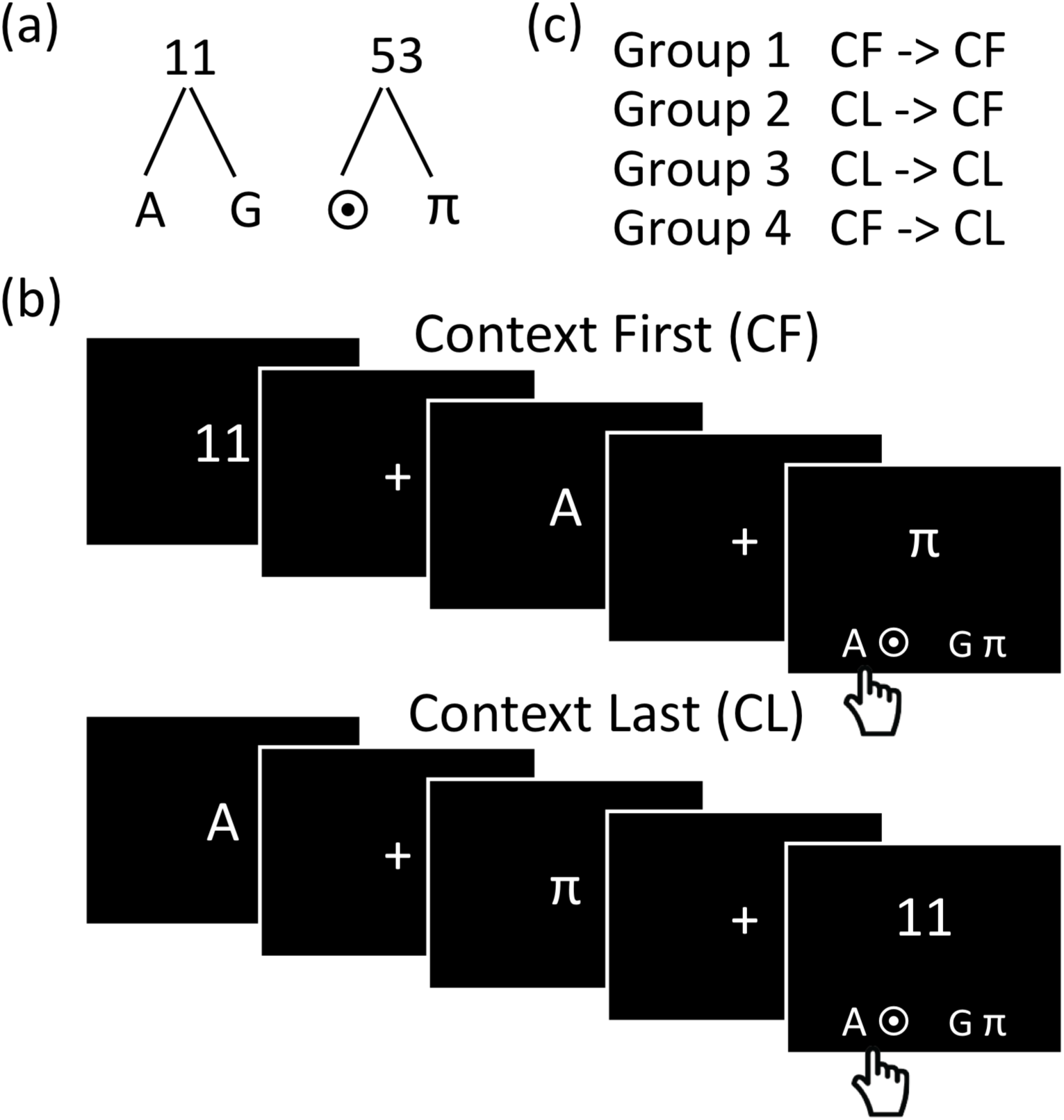
Experiment 1 task and experiment design. (a) Rule trees linking contextual items to lower-level items. Participants memorized the rule trees prior to beginning the task and were given an opportunity to review at the beginning of each block. (b) Sample context first (CF) and context last (CL) trials. The contextual item (number) indicated which of the other two items in the sequence is the target item. Each trial concluded with a response panel that consisted of two pairs of items. Participants indicated with a key press which pair contained the correct target item (denoted by a hand with a tick mark). (C) Participants were assigned to one of four experimental groups. Each group completed two blocks (‘training’ and ‘test’), whose trial structure was determined by group membership.

To illustrate a trial, consider the sequence 11…A…π (Figure 2(b)). As per the rule trees, 11 indicates that either of the letters, A or G, would be a target on that trial. Therefore, in the above example sequence, A, and not π, is the target. Each trial concluded with a response panel (presented at the same time as the last item in the sequence) that consisted of two pairs of items at the lower left or right of the screen.

Participants had to indicate whether the target item for that trial appeared as part of the left pair (left key) or the right pair (right key). In the example, the left key would be the correct response as it contained the ‘A’ target. The location of the letters and symbols in the target panel was randomized trial-to-trial such that all items appeared with equal frequency on the left and right across all trials. Further, 50% of the trials were ‘incongruent’ in that the lower-level items which appeared in the sequence were associated with different response keys in the response panel, and participants could not perform accurately without attending to the contextual cue. Participants were instructed to respond as fast as possible, while being accurate. Response panels were randomized so that on half the trials the left key was the correct response.

Apart from the position of the contextual item in the sequence (either first or last), the position of the lower level items was randomized and balanced between first, last, or middle position in the three-item sequence, such that there were equal numbers of each possible trial type in the block. All stimuli were presented in white, on a black background. The first two items in the sequence were each presented for 300 ms, while the final item (along with the response panel) was self-timed with response required within a window of 3000 ms. The inter-stimulus interval was jittered between 600 ms and 1600 ms, while the inter-trial interval was fixed at 500 ms. A fixation cross was displayed during each interval.

Two different trial structures were employed (Figure 2(b)). In the ‘context first’ (CF) structure, the contextual cue (the number) appeared at the beginning of the sequence. In the ‘context last’ (CL) structure, the context cue appeared at the end of the sequence. CF and CL trial structures afforded the use of different WM gating policies: CL trials required a ‘selective output-gating policy’, whereas efficient behavior in CF trials could be supported by a ‘selective input-gating policy’. Instructions were worded to be neutral with respect to trial-structure and examples of both trial structures were provided to participants during instruction (see Appendix A). They were also explicitly told that they might see both kinds of trial structures during the experiment. Participants were not given any prior practice on either trial structure.

Participants were randomly assigned to one of four groups. Each group carried out two, forty-eight trial blocks of the task: a ‘training’ block and a ‘test’ block (Figure 2(c)). Within each block, the trial structure (CF or CL) was always fully predictable, providing an opportunity to learn an efficient gating policy. For two groups, the trial structure remained the same between blocks (CF -> CF or CL -> CL). For the other two groups, the trial structure switched (CL -> CF or CF -> CL). Importantly, the task rules (Figure 2(a)) always remained the same across blocks. Therefore, the design allowed us to dissociate transfer effects from any effects of rule learning.

We subsequently ran two additional group of participants. One group (CF -> CL -> CF), was similar to the CL -> CF group in that they carried out a CF test block immediately after a CL training block. But this group also carried out a block of CF trials prior to the CL block. This allowed us to test the possibility that performance decrements observed in the CL -> CF group test block were simply due to the obligatory perseveration of the last-used gating policy rather than a deliberate transfer of a gating policy. Participants in this group had the chance to learn both selective-input gating and selective-output gating policies prior to the CF test block. An obligatory perseveration account would predict that the CL -> CF group and the CF -> CL -> CF group participants would show similar initial performance in the test block. On the other hand, a deliberate transfer account would predict that the CF -> CL -> CF group would adapt more quickly to the CF test block thus showing improved initial performance.

Another group (CF2 -> CF) served to control for the effects of WM load in the CL -> CF group. In the training block of the CL -> CF group, two items must be maintained in WM memory on each trial, making the task more difficult. Therefore, any negative transfer effects observed may be due to the fatigue or motivation loss induced by the higher load or the difficulty of the task rather than the change in trial structure. To control for this, in the CF2 -> CF group, the training task consisted of rules in which each contextual item was associated context with four lower-level items. On each trial, context was presented first, followed by two pairs of items, one of which was relevant. Participants were required to keep both relevant lower-level items in WM, as either of them could be the target item. Therefore, the CF2 -> CF served as a control with equivalent WM load during training as the CL -> CF group, but no change in trial structure.

##### Data Analysis

Analyses utilized response accuracy and response time (RT) measures. All RT analyses focused only on trials where participants responded correctly. In addition, we discarded trials in which participants responded in less than 250 ms. RTs were log-transformed prior to statistical analyses. For analyses of transfer effects, we computed mean RT and response accuracy separately for eight trial bins of six trials each, motivated by our prediction that transfer would most affect the initial trials in a task.

### Results

Overall, participants performed well at the task (Table 1). As seen in previous studies (Chatham et al., 2014), CF was performed more accurately (94.4% vs. 90.9%; [*t* (112) = 2.38, *p* = 0.019, *d* = 0.44]) and more efficiently (915 ms vs. 1261 ms; [*t* (112) = 7.94, *p* < 0.001, *d* = 1.19]) than CL, providing evidence that participants take advantage of the trial structure to selectively input gate lower level items. Note that this was despite the fact that the context stimulus was displayed for longer in the CL condition than in the CF condition. Summary statistics of performance for each group and each block are shown in Table 2.

**Table 1:** Experiment 1 & 2 summary of performance data^a^

| *Trial structure* | *Accuracy*<br>*(% correct)* | *Response time*<br>*(ms)* |
| --- | --- | --- |
| *Experiment 1* |  |  |
| CF | 94.4 ± 0.02 | 915 ± 60 |
| CL | 90.9 ± 0.02 | 1261 ± 61 |
| Overall | 92.6 ± 0.01 | 1088 ± 53 |
| *Experiment 2* |  |  |
| CF | 92.2 ± 0.09 | 881 ± 294 |
| CL | 90.6 ± 0.08 | 1250 ± 236 |
| Overall | 91.4 ± 0.08 | 1066 ± 267 |

<sup>a</sup>Summary statistics are means and 95% confidential intervals.

**Table 2:** Experiment 1 & 2 group-wise summary of performance data^a^

| *Group* | *Training block(s)* |  | *Test block* |  |
| --- | --- | --- | --- | --- |
|  | *Accuracy*<br>*(% correct)* | *Response time*<br>*(ms)* | *Accuracy*<br>*(% correct)* | *Response time*<br>*(ms)* |
| *Experiment 1* |  |  |  |  |
| CF → CF | 92.0 ± 0.05 | 971 ± 118 | 97.4 ± 0.01 | 907 ± 94 |
| CL → CF | 91.6 ± 0.04 | 1304 ± 102 | 95.3 ± 0.03 | 910 ± 96 |
| CL → CL | 88.3 ± 0.06 | 1317 ± 92 | 94.5 ± 0.02 | 1266 ± 83 |
| CF → CL | 92.7 ± 0.03 | 896 ± 118 | 89.7 ± 0.04 | 1188 ± 125 |
| CL2 → CF | 92.6 ± 0.06 | 1182 ± 103 | 94.5 ± 0.09 | 806 ± 92 |
| CF → CL → CF | CF: 91.8 ± 0.06<br>CL: 90.4 ± 0.04 | CF: 889 ± 111<br>CL: 1176 ± 84 | 92.5 ± 0.04 | 741 ± 60 |
| *Experiment 2* |  |  |  |  |
| *Same rules for each block* |  |  |  |  |
| CF → CF | 94.2 ± 0.03 | 905 ± 116 | 96.1 ± 0.04 | 839 ± 98 |
| CL → CF | 93.8 ± 0.06 | 1299 ± 135 | 90.1 ± 0.05 | 883 ± 162 |
| CL → CL | 94.1 ± 0.02 | 1175 ± 77 | 95.0 ± 0.03 | 1208 ± 71 |
| CF → CL | 87.2 ± 0.02 | 928 ± 182 | 90.5 ± 0.03 | 1367 ± 121 |
| *Different rules for each block* |  |  |  |  |
| CF → CF | 95.4 ± 0.02 | 820 ± 69 | 95.0 ± 0.02 | 806 ± 65 |
| CL → CF | 89.8 ± 0.05 | 1202 ± 110 | 87.2 ± 0.07 | 983 ± 174 |

<sup>a</sup>Summary statistics are means and 95% confidence intervals.*Transfer effects.*

#### Transfer effects

Our primary question is whether working memory gating policies learned in one block are transferred to a subsequent block. Such transfer predicts improvements (positive transfer) or decrements (negative transfer) in performance on subsequent blocks conditional on the nature of the prior experience. To this end, we focus on the effects of prior experience with a trial structure on performance with the same or new trial structure in subsequent blocks. A summary of the ANOVAs performed to test for transfer effects are summarized in Table 3.

**Table 3:** Experiment 1 summary ANOVA results

| *Statistical test* | *Effect* | *df* | *F* | *p* | $\eta_p^2$ |
| --- | --- | --- | --- | --- | --- |
| *Negative transfer Effects* |  |  |  |  |  |
| CF trials response time:<br>Trial Bin x Training History<br>mixed-model ANOVA | Trial bin ME* | (7, 252) | 7.82 | <0.001 | 0.178 |
|  | Training history ME | (1, 36) | 0.02 | >0.25 | 0.001 |
|  | Trial bin x training<br>history interaction* | (7, 252) | 3.15 | 0.003 | 0.081 |
| CF trials response accuracy:<br>Trial Bin x Training History<br>mixed-model ANOVA | Trial bin ME | (7, 252) | 1.73 | 0.103 | 0.046 |
|  | Training history ME | (1, 36) | 1.27 | >0.250 | 0.034 |
|  | Trial bin x training<br>history interaction | (7, 252) | 1.12 | >0.250 | 0.030 |
| CL trials response time:<br>Trial Bin x Training History<br>mixed-model ANOVA | Trial bin ME* | (7, 252) | 3.13 | 0.003 | 0.080 |
|  | Training history ME | (1, 36) | 1.92 | 0.174 | 0.051 |
|  | Trial bin x training<br>history interaction | (7, 252) | 0.943 | >0.250 | 0.026 |
| CL trials response accuracy:<br>Trial Bin x Training History<br>mixed-model ANOVA | Trial bin ME | (7, 252) | 2.36 | 0.065 | 0.051 |
|  | Training history ME | (1, 36) | 0.16 | 0.249 | 0.004 |
|  | Trial bin x training<br>history interaction* | (7, 252) | 2.71 | 0.03 | 0.040 |
| *Positive transfer Effects* |  |  |  |  |  |
| CF trials response time:<br>Trial Bin x Block repeated-<br>measures ANOVA | Trial bin ME | (7, 126) | 1.04 | >0.250 | 0.055 |
|  | Block ME * | (1, 18) | 4.55 | 0.047 | 0.202 |
|  | Trial bin x block* | (7, 126) | 3.64 | 0.001 | 0.168 |
| CF trials response accuracy:<br>Trial bin x block repeated-<br>measures ANOVA | Trial bin ME* | (7, 126) | 1.21 | >0.250 | 0.063 |
|  | Block ME | (1, 18) | 4.16 | 0.056 | 0.188 |
|  | Trial bin x block | (7, 126) | 0.55 | >0.250 | 0.030 |
| CL trials response time:<br>Trial bin x Block repeated-<br>measures ANOVA | Trial bin ME* | (7, 126) | 2.62 | 0.030<sup>+</sup> | 0.127 |
|  | Block ME | (1, 18) | 1.78 | 0.202 | 0.089 |
|  | Trial bin x block | (7, 126) | 1.04 | >0.250 | 0.054 |
| CL trials response accuracy:<br>Trial bin x Block repeated-<br>measures ANOVA | Trial bin ME | (7, 126) | 3.27 | 0.053 | 0.193 |
|  | Block ME | (1, 18) | 4.30 | 0.012<sup>+</sup> | 0.154 |
|  | Trial bin x block | (7, 126) | 1.72 | 0.127<sup>+</sup> | 0.067 |
| *Target position effects* |  |  |  |  |  |
| CF trials response time:<br>Trial bin x Training history<br>mixed-model ANOVA | Trial bin ME* | (1, 36) | 6.01 | 0.019 | 0.143 |
|  | Training history ME* | (1, 36) | 0.07 | >0.250 | 0.001 |
|  | Trial bin x training<br>history interaction* | (1, 36) | 4.55 | 0.039 | 0.114 |
| CL trials response time:<br>Trial bin x Training history<br>mixed-model ANOVA | Trial bin ME | (1, 36) | 0.06 | 0.068 | 0.002 |
|  | Training history ME* | (1, 36) | 0.49 | >0.250 | 0.014 |
|  | Trial bin x training<br>history interaction* | (1, 36) | 1.05 | >0.250 | 0.029 |

First, we compared performance data from *test blocks* when the trial structure changed, CL -> CF and CF -> CL, with the corresponding *training blocks* with the same trial structure. Log-transformed RT data from the blocks where the structure changed were analysed using two separate mixed-model ANOVAs (for CF and CL trial structures) with trial bin (1-8) as a within-subject factor and training history as a between-subject factor. Note that the training history factor has two levels – ‘no prior training’ and ‘prior training with different structure’. This reflects a comparison between the training block in the no-change group (CF -> CF, for example) and the test block in the structure-change group (CL -> CF, for example)).

For the CF condition (Figure 3(a)), the ANOVA revealed a main effect of trial bin [*F* (7, 252) = 7.82, *p* < 0.001, η_p_ ^2^ = 0.178] and a trial bin x training history interaction [*F* (7, 252) = 3.15, *p* = 0.003, η_p_ ^2^ = 0.081]. A post-hoc *t*-test [*t* (36) = 2.75, *p* = 0.009, *d* = 0.89] showed a trend of the participants in the CL -> CF being slower in the 1^st^ trial bin though this difference was marginal after a Bonferroni correction.

**Figure 3:**
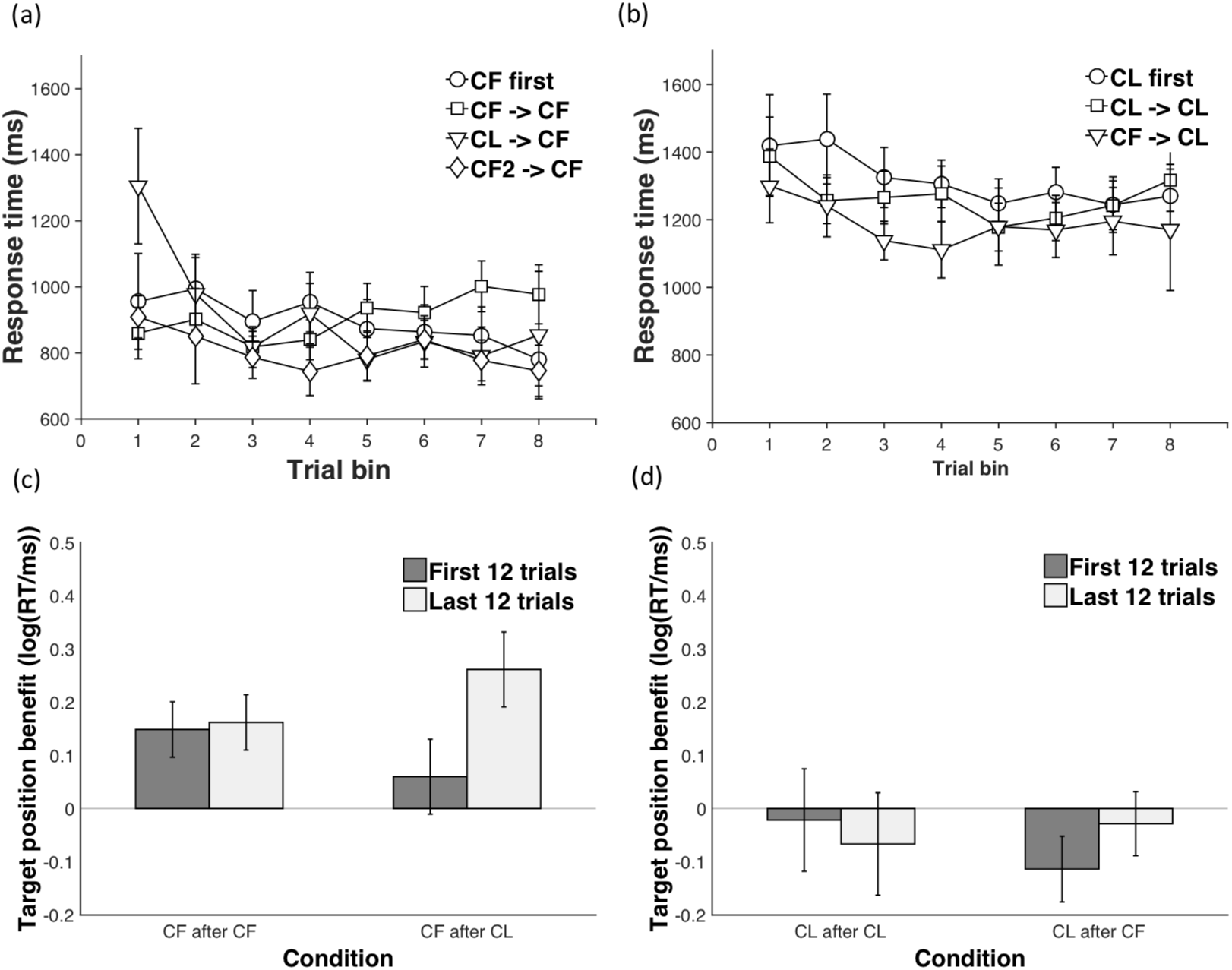
(a) and (b) - Transfer effects for CF (a) and CL (b) trial structures. Mean RT for each 6-trial bin is plotted as a function of training history - no prior training (circles), prior training with same structure (squares), or different trial structure (triangles). Error bars reflect 95% confidence intervals. (c) and (d) - Target position effects for CF (c) and CL (d) trial structures. Mean target position benefit in RT (target-middle RT – target-last RT) plotted for early (first 12 trials – dark grey bars) and late (last 12 trials – light grey bars) in test blocks of CF -> CF and CL -> CF conditions (c) and CL -> CL and CF -> CL conditions (d). RTs were log-transformed. Error bars reflect 95% confidence intervals.

It is possible that the effect of prior training with the CL trial structure on CF response times was driven by differences in working memory load (CL trial structures produce a higher WM load) or task difficulty during training. To rule out this possibility, we considered performance of the CF2 -> CF group. In the CF2 training block, context was always presented first, but participants were required to maintain two lower-level items in WM rather than just one (see methods). Therefore, the task produced an equivalent WM load during training as the CL -> CF group, but no change in trial structure. Indeed, response times in the CF2 training block were significantly higher than in a CF training block [*t*(36) = 3.72, *p* < 0.001, *d* = 0.98], and somewhat lower than the CL training block [two-sample *t-*test: *t*(36) = -2.2 *p* = 0.054, *d* =0.80]. To test whether the slower responding observed in the test block of the CL -> CF group could be accounted for by WM load or difficulty, we first ran a mixed-model ANOVA comparing the test block of the CF2 -> CF group with the training block of the CF -> CF group (analogous to the earlier comparison of the CL -> CF test block with the CF -> CF test block). If the higher WM load contributes to slowing in the test block, we should observe an analogous trial bin x training history interaction. However, while we found a significant main effect of trial bin [*F* (7, 252) = 2.68, *p* = 0.024 (Greenhouse-Geisser corrected), η_p_ ^2^ = 0.069], there was neither a significant main effect of the training history [*F* (1, 36) = 1.21, *p* > 0.1, η_p_ ^2^ = 0.032], nor a significant trial bin x training history interaction [*F* (7, 252) = 1.31, *p* > 0.1, η_p_ ^2^ = 0.035]. To confirm that participants showed a slowing in responses in the CL -> CF test block over and above that caused by the WM load, we ran another mixed-model ANOVA, this time directly comparing the test blocks of the CF2 -> CF and CL -> CF groups. We found a significant trial bin x training history interaction [*F* (7, 252) = 2.78, *p* = 0.008, η_p_ ^2^ = 0.072] along with a significant main effect of trial bin [*F* (7, 252) = 8.91, *p* < 0.001, η_p_ ^2^ = 0.198].

For the CL condition (Figure 3(b)), the main effect of trial bin was significant [*F* (7, 252) = 3.13, *p* = 0.003, η_p_ ^2^ = 0.08] reflecting a quadratic speeding or RT over trial bin [*F*(1, 36) =14.02, *p* = 0.001, η_p_ ^2^ = 0.280], but the trial bin x training history interaction was non-significant [*F*(7, 252) = 0.94, *p* > 0.250, η_p_ ^2^ = 0.026].

Similar analyses of accuracy data for the CF and CL conditions are summarized in Table 3. In summary, we found no evidence for performance decrements. We did observe an improvement in accuracy in the CF -> CL condition test block, compared to the CL -> CL training block as evidenced by a training history x trial bin interaction. Post-hoc tests comparing each trial bin were non-significant after a Bonferroni correction. Two participants in the CL -> CL group performed poorly in the training block (< 60% correct) and may be driving this improvement. The ANOVA after removing these two outliers again showed a significant, but less reliable trial bin x training history interaction [*F* (1, 16) = 2.39, Greenhouse-Geisser corrected *p* = 0.044, η_p_ ^2^ = 0.07].

Next, we compared the training and test blocks RTs from participants in the CF -> CF and CL -> CL groups using a trial bin x block ANOVA. A transfer of a WM gating policy would predict improved performance (positive transfer). Data from the CF -> CF group were analysed with a repeated measures ANOVA with trial bin (8 levels: 1-8) and block (2 levels: training or test) as within subject factors. The main effect of trial bin was non-significant [*F* (7, 126) = 1.04, *p* > 0.250, η_p_ ^2^ = 0.055], while the main effect of block was marginal [*F* (1, 18) = 4.55, *p* = 0.047, η_p_ ^2^ = 0.202]. On the other hand, the trial bin x block interaction was significant [*F* (7, 126) = 3.64, *p* = 0.001, η_p_ ^2^ = 0.168]. Bonferroni-corrected (*α* = 0.006), post-hoc paired *t*-tests revealed that this interaction was driven by faster RTs in the first trial bin in the test block compared to the training block [*t* (18) = 3.31, *p* = 0.004, *d* = 0.76]. A similar analysis of the CL -> CL group data revealed a main effect of trial bin [*F*(7, 126) = 2.62, *p* = 0.015, η_p_ ^2^ = 0.127], while the main effect of block [*F*(7, 126) = 1.76, *p* = 0.100, η_p_ ^2^ = 0.089], and the trial bin x block interaction were non-significant [*F*(7, 126) = 1.04, *p* > 0.250, η_p_ ^2^ = 0.054].

ANOVA of the accuracy data in the CF -> CF group showed a marginal main effect of block [*F* (1, 18) = 4.56, *p* = 0.056, η_p_ ^2^ = 0.188]. All other effects were non-significant [*p* > 0.1]. For the CL -> CL group, again, the main effect of block was marginal [*F* (1, 18) = 4.3, *p* = 0.053, η_p_ ^2^ = 0.193] and the main effect of trial bin was significant [*F* (7, 126) = 3.27, *p* = 0.003, η_p_ ^2^ = 0.54]. The interaction was non-significant [*F* (7, 126) = 1.83, *p* = 0.088, η_p_ ^2^ = 0.09]. Two participants in the CL -> CL group performed poorly in the training block (< 60% correct). The ANOVA after removing these two outliers showed no main effect of block [*F* (1, 16) = 1.89, *p* > 0.1, η_p_ ^2^ = 0.11], a marginal effect of trial bin [*F* (7, 112) = 2.13, *p* = 0.05, η_p_ ^2^ = 0.12] and a significant marginal block x trial bin interaction [*F* (7, 112) = 2.28, *p* = 0.033, η_p_ ^2^ = 0.125] with a trend toward improved accuracies in the first trial bin [*t* (16) = 2.46, *p* = 0.025, *d* = 0.60] that did not survive a Bonferroni correction.

In summary, we found evidence of asymmetric negative transfer effects. Participants with prior experience with the CL trial structure responded more slowly on early trials of a subsequent CF block compared to those who had no prior experience. On the other hand, prior training with the CF trial structure had no effect on performance in a subsequent CL block. We also found evidence of performance improvements on repetition of the same trial structure. Participants with prior experience with the CF trial structure were significantly faster on the early trials of a subsequent CF block. Participants in the CF -> CF groups were also marginally more accurate in the test block compared to the training block, while those in the CL -> CL group showed a trend toward more accurate performance in the first trial bin of the test block. These performance improvements are consistent with the positive transfer of a gating policy, though they may also reflect practice with the S-R rules.

##### Obligatory v/s deliberate transfer

In the previous analyses, we found that participants with prior experience with the CL trial structure responded more slowly on early trials of a subsequent CF block compared to those who had no prior experience. An important question concerns whether this effect was due to participants deliberately transferring a gating policy, or simply perseverating with the previously used policy in an obligatory manner. To test this, we compared the test blocks from the CF -> CL -> CF group and the CL -> CF groups in a mixed-model ANOVA with trial bin and training history as factors. If participants briefly perseverate with the previously used policy, we would expect participants in the CF -> CL -> CF group to show some slowing in RT as the CL -> CF group. On the other hand, if transfer is deliberate, the CF -> CL -> CF group would be expected to reinstate the selective input gating policy they learnt in the initial CF block, thus producing little or no slowing. We found a significant main effect of trial bin [*F* (7, 252) = 9.28, *p* < 0.001, η_p_ ^2^ = 0.205] and a significant trial bin x training history interaction [*F* (7, 252) = 4.3, *p* = 0.003, η_p_ ^2^ = 0.080]. Participants in the CL -> CF group were indeed slower in the first and second trial bin of the test block compared to participants in the CF -> CL -> CF group as confirmed by Bonferroni-corrected t-tests [Trial bin 1: *t* (18) = 3.68, *p* < 0.001, *d* = 1.03; Trial bin 2: *t* (18) = 3.35, *p* = 0.002, *d* = 0.96].

Comparison of the test blocks in the CF -> CL -> CF group with the training block of the CF -> CF group (just as for the CL -> CF group in the prior section) showed a marginally significant main effect of trial bin [*F* (7, 252) = 2.11, *p* = 0.04, η_p_ ^2^ = 0.05] and training history [*F* (1, 36) = 1.98, *p* = 0.038, η_p_ ^2^ = 0.205] and a significant trial bin x training history interaction [*F* (7, 252) = 2.33, *p* = 0.026, η_p_ ^2^ = 0.061]. But the main effect of training history and the train bin x training history interaction were driven by *faster responses* in the CF -> CL -> CF test block compared to the CF -> CF training block. Therefore, despite the immediately previous CL block, participants were in fact faster compared to a CF training block, presumably due to a combination of positive transfer of a gating policy learnt in the earlier CF block, and practice with the S-R mappings.

To further test the unique effects of and minimize the contribution of practice, we compared the RTs in the first trial bin of the CL -> CF group and the CF -> CL -> CF while correcting for baseline differences (respectively, the CF -> CF training and test blocks) in a general linear model. We evaluated the contrast [(CL -> CF test) – (CF -> CF training)] – [(CF -> CL -> CF test) – (CF -> CF test)] and found a significant effect [*F* (7, 252) = 5.44, *p* = 0.023, η_p_ ^2^ = 0.633]. This result makes it unlikely that participants simply perseverated with the last used policy in an obligatory manner. Instead, our results are consistent with a deliberate transfer of a selective output gating policy.

##### Target position effects

In the previous section, we found evidence consistent with transfer of gating policies across task contexts. On this account, participants learn a gating policy in the training block which they transfer to the test block. Initial performance in the test block, is supported by the transferred policy. Participants in the CF -> CF group learn a selective input-gating policy in the initial CF training block thus enabling efficient performance in the subsequent CF test block. Participants in the CL -> CF group, on the other hand, learn a selective output-gating policy in the training block. This policy continues to support performance in the changed circumstances of the CF test block, but at the cost of slower responses. In contrast, in the CF -> CL group, a transferred selective input-gating policy would not support performance in the changed circumstances, thus immediately requiring the deployment of a new policy. This account predicts that initial behaviour in the CL -> CF test block is under the control of a selective output-gating policy, while initial behaviour in the CF -> CF test block is under the control of a selective input-gating policy, despite both the block being identical.

A feature of the task design allows us to test these predictions. CF trials come in two forms, those in which the target item is in the second position (target middle), and those in which the target item is in the last position (target last). The final response decision requires knowledge of the target item, and therefore cannot be initiated or prepared for until target selection has taken place. Critically, the timing of target selection is contingent on the kind of gating policy that one employs. Under a selective output-gating policy, target selection does not proceed until the final item is presented. Therefore, no benefit can be gained on target-middle trials on which target information is available early. On the other hand, under a selective input-gating policy, target selection occurs at the level of input-gating. Therefore, on target-middle CF trials, target selection can occur once the second item in the sequence is processed, and one can begin preparing for the upcoming response decision; for example, setting up an attentional set to search for the target item in the response panel. Thus, under an input-gating policy only, a *target position benefit* can be expected on target-middle vs. target-last trials. No such benefit would follow from an output-gating policy.

Consistent with the account developed above, if participants in the CL -> CF group transfer a selective output-gating policy from the training block, target position benefits should be unavailable on the early trials of the test block. As participants learn a selective input-gating policy with experience, target position benefits would become available later in the block. On the other hand, in the CF -> CF test blocks participants are more likely to transfer a selective input-gating policy (Chatham et al., 2014) and robust target position benefits should be observed on the early trials as well. Finally, target position benefits should not be observed at all in the CL -> CL and CF -> CL test blocks, which thus serve as a negative control.

Target position benefits were computed by subtracting log-transformed RTs on target-middle trials from those on target-last trials and are plotted in Figure 3 ((c) & (d)). A summary of the ANOVAs performed to test for transfer effects are summarized in Table 3. To begin with, we employed one-sample *t*-tests to examine the presence of target position benefits. As predicted, no target position benefit was found on the early (first 12) trials of the test block in the CL -> CF group [one sample *t* (18) = 0.89, *p* > 0.250, *d* = 0.2]. On the other hand, a robust target position effect was observed on the late (last 12) trials [one-sample *t* (18) = 5.62, *p* < 0.001, *d* = 1.29]. A paired *t*-test confirmed a significant effect of trial bin on the size of the target position benefit [*t* (18) = 2.86, *p* = 0.010, *d* = 0.66].

We next compared target position benefits in the CL -> CF and CF -> CF test blocks. A mixed-model ANOVA with trial bin (2 levels: first 12 trials bin, last 12 trials bin) as a within-subject factor and group (2 levels: CF -> CF, CL -> CF) as a between-subject factor confirmed a significant trial bin x group interaction [*F* (1, 36) = 4.61, *p* = 0.039, η_p_ ^2^ = 0.114]. As a final control, we examined target position benefits in the CL -> CL test block and found no evidence of a target position benefit. On the early trials, however, there was a slight target position cost [*t* (18) = -2.6, *p* = 0.020, *d* = -0.6].

In summary, these results provide specific and clear evidence for the transfer of a selective output-gating policy from the training to the test block in the CL -> CF group, and the learning of selective input-gating policy with experience in the test block.

## Experiment 2

The results of Experiment 1 provide evidence for the transfer of working memory gating policies. We observed an asymmetric negative transfer across changes in trial structure, even as SR rule structure was held constant. We also observed target-position effects that provided specific evidence for the transfer of a selective output-gating policy learnt in the CL task context to a CF task context. In addition, we found an improvement in performance when the same trial structure was held constant across the training and the test blocks. Such improvements are consistent with the transfer of a gating policy, but could also be explained with a simpler practice account because the S-R rules were also held constant.

In Experiment 2, we attempted to replicate and extend these findings. We carried out a stronger test of the hypothesis that gating policies and S-R rules are separable forms of task knowledge by independently manipulating prior experience with an S-R rule vs. a trial structure. This also enabled to test between the transfer and practice accounts of the performance improvements. We replicated the effects of prior experience with a trial structure, and found that changing the rules has little effect on later performance.

### Methods

#### Participants

117 adult, right-handed participants (45 males, 72 females; age-range: 18-30, M = 21.6, SD = 2.6) from the Providence, RI area were recruited to take part in the computer-based experiment. We endeavored to collect between 18-20 participants in each of six groups based on approximate effect sizes suggested by pilot data and Experiment 1. 4 participants were excluded as they were on psychoactive medication. 5 participants were excluded because of low performance (< 70% accuracy) on the task. This left 108 participants (40 males, 68 females; age-range: 18-30, M = 21.4, SD = 2.6). All remaining participants had normal or corrected-to-normal vision, and no reported neurological or psychological disorders. All participants gave informed, written consent as approved by the Human Research Protections Office of Brown University, and they were compensated for their participation.

#### Task and Experiment Design

Experiment 2 used the same task as Experiment 1. Participants were randomly assigned to one of six experimental groups. Four of these groups performed identical transfer conditions to the four conditions used in Experiment 1. Two additional groups (CF -> CL (different rules) and CL -> CF (different rules), were presented with novel rules at the beginning of each block. The new rules utilized the same categories (numbers for context, letters and symbols for lower-level items) but a novel set of numbers, letters and symbols. The assignment of each rule set to the training and test block was counterbalanced across participants and groups.

### Results

Overall performance was strong and comparable between the two experiments (Table 1). Summary statistics for all measures are displayed in Table 2. Results are presented in two sections. In the first section, we replicate the transfer effects and target position effects from Experiment 1. In the final section, we turn to a comparison of the same-rule and different-rule groups.

#### Transfer effects and target position effects

Behavioural results from the same rule conditions are plotted in Figure 4 and the results of the ANOVAs are presented in Table 4. In summary, we replicated the asymmetric negative transfer effects from Experiment 1, finding a significant trial bin x training history interaction [*F* (7, 238) = 4.99, *p* < 0.001, η_p_ ^2^ = 0.128] on RTs in the CF condition (Figure 4(a)), but not the CL condition (Figure 4(b)). We also obtained an asymmetric positive transfer effect, with a significant trial bin x block interaction on response accuracies in the CF -> CF, [*F* (7, 238) = 4.90, *p* < 0.001, η_p_ ^2^ = 0.214], but not the CL -> CL conditions.

**Table 4:**
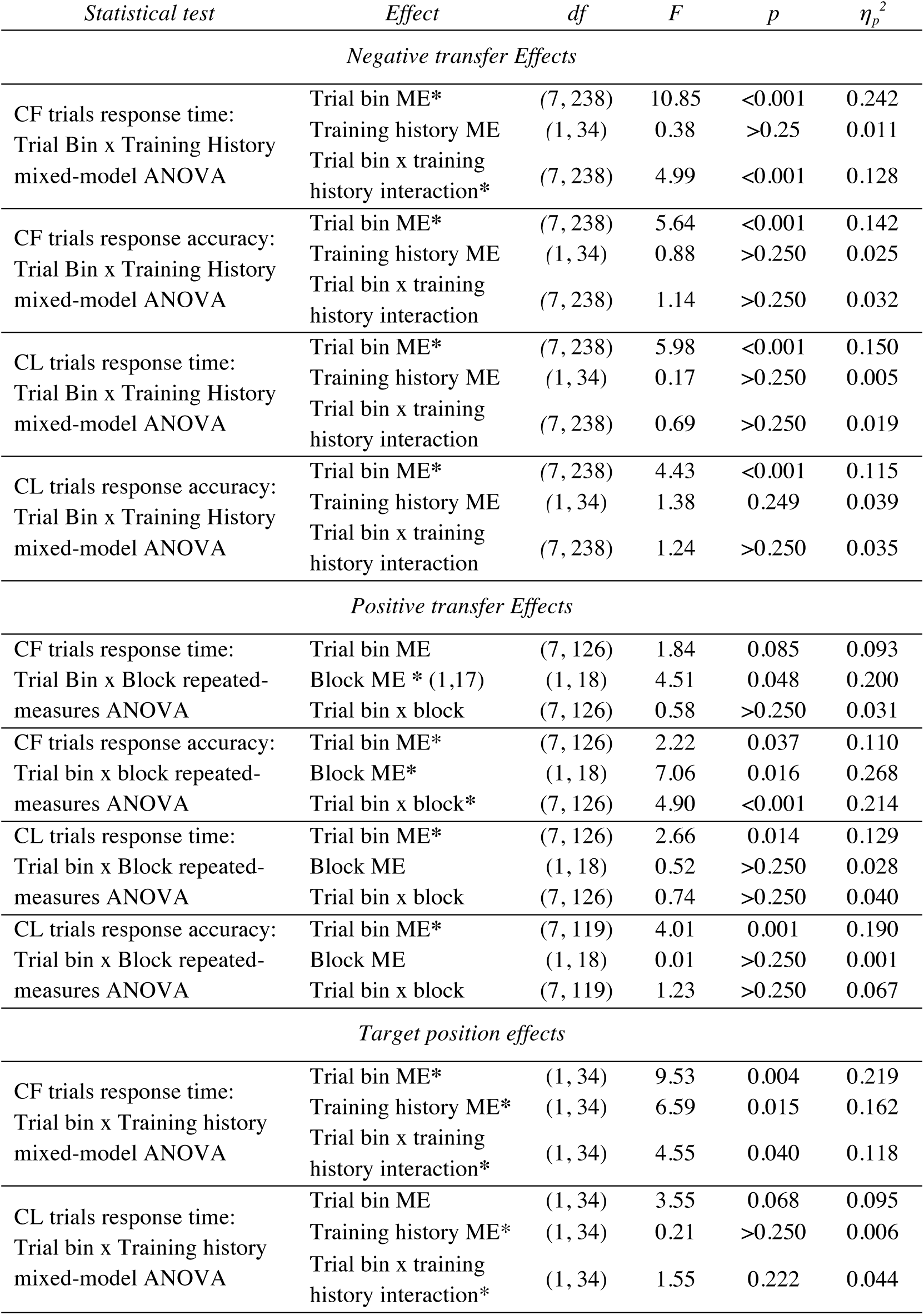
Experiment 2 summary ANOVA results

**Figure 4:**
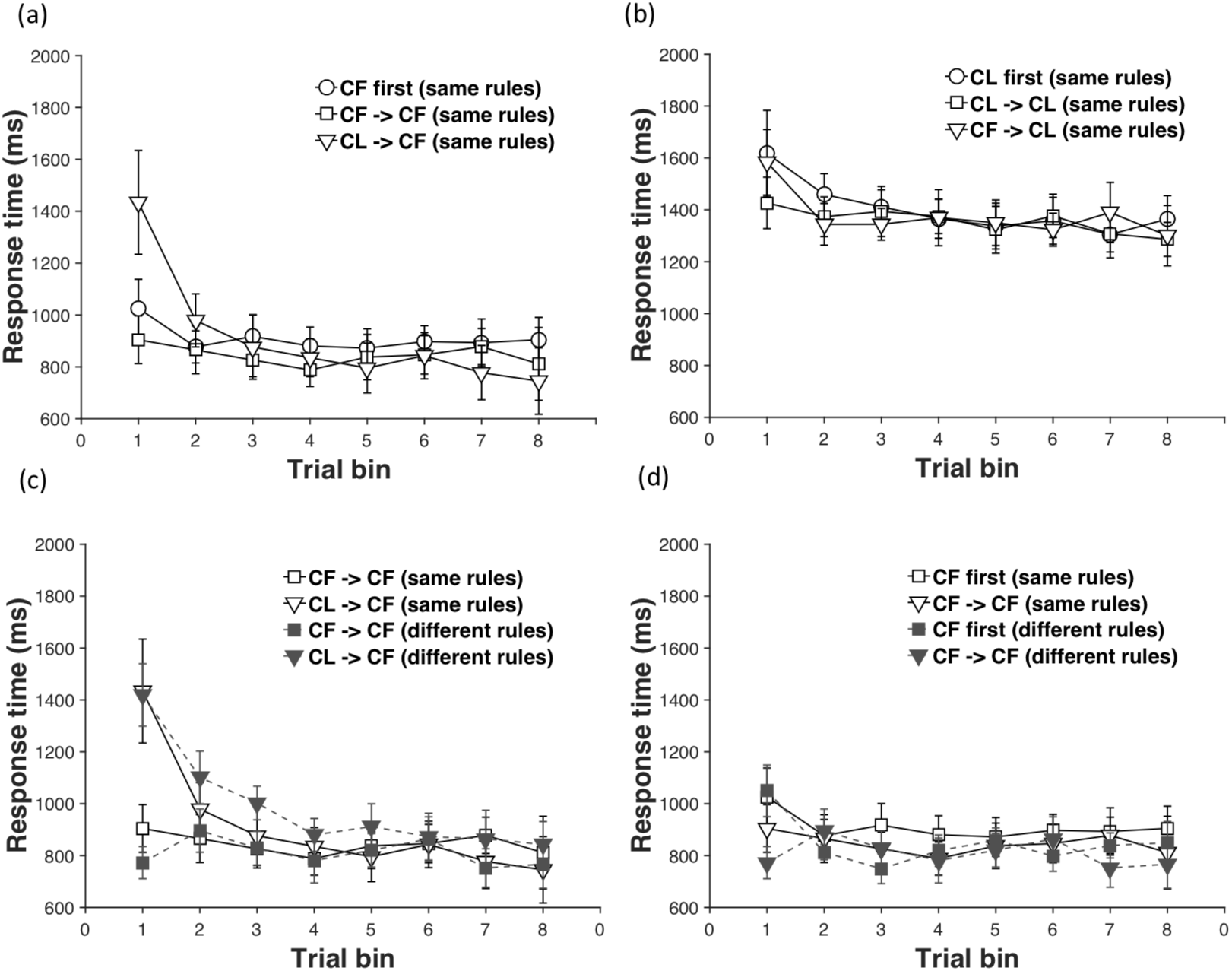
Results from Experiment 2. (a) & (b) – Transfer effects for CF (a) and CL (b) trial structures when rules remain the same. Mean RT for each 6-trial bin is plotted as a function of training history - no prior training (circles), prior training with same structure (squares), or different trial structure (triangles). Transfer effects observed in Experiment 1 are replicated here. (c) & (d) - Comparison of negative (c) and positive (d) transfer effects in the different-rule (open shapes, dashed lines) and corresponding same-rule groups (filled shapes, solid lines). The transfer effects are primarily driven by previous experience with a trial structure rather than with a rule. Error bars reflect 95% confidence intervals.

We also replicated the pattern of target position effects observed in Experiment 1. A non-significant trend for a target position cost was observed on the early (first 12) trials of the test block in the CL -> CF group [one sample *t* (17) = -1.92, *p* = 0.072, *d* = -0.45]. A significant target position benefit was observed on the late (last 12) trials [one-sample *t* (17) = 2.83, *p* = 0.012, *d* = 0.67]. A paired *t*-test confirmed a significant effect of trial bin on the size of the target position benefit [*t* (18) = 2.95, *p* = 0.009, *d* = 0.70]. ANOVAs (full detail provided in Table 4) confirmed that a significant trial bin (first 12 vs. last 12 trials) x training history interaction was present for the CF condition, [*F* (7, 238) = 4.55, *p* = 0.04, η_p_ ^2^ = 0.118] but not for the CL condition [*F* (7, 238) = 1.55, *p* = 0.222, η_p_ ^2^ = 0.044].

#### Effect of rule versus trial structure change

Finally, we turn to a comparison of the same-rule and different-rule groups and examine the relative influence of prior experience with rules vs. trial structures on subsequent performance.

Log-transformed RT data from the test blocks of four groups: CF -> CF (same rule), CL -> CF (same rule), CF -> CF (different rule) and CL -> CF (different rule) (Figure 4(c) and 4(d)) were analysed with a mixed-model with trial bin (8 levels: 1-8) as a within-subject factor and previous trial structure (2 levels: same trial structure, different trial structure) and previous rule (2 levels: same rule, different rule) as between-subject factors. The ANOVA revealed a significant main effect of trial bin [*F* (7, 476) = 19.39, *p* < 0.001, η_p_ ^2^ = 0.222] and a trial bin x previous trial structure interaction [*F* (7, 476) = 13.31, *p* < 0.001, η_p_ ^2^ = 0.164]. Importantly, neither the main effect of the previous rule [*F* (7, 476) = 0.001, *p* > 0.250, η_p_ ^2^ = 0.00], nor the three-way interaction between trial bin, previous trial structure, and previous rule [*F* (1, 68) = 0.03, *p* > 0.250, η_p_ ^2^ = 0.00] was significant, suggesting that prior experience with a rule had no influence on subsequent performance. To further confirm that prior experience with a rule did not subtly influence the slope of the RT curves in the CL -> CF test blocks (Figure 4(c)), we fit power functions to each subject’s curve and entered the estimated parameters into an independent-sample *t*-test comparing the rule change and no rule change group. We found no reliable differences [*t* (34) = 0.68, *p* = 0.208, *d* = 0.23].

Analysis of the accuracy data, similarly, revealed a significant main effect of previous trial structure [*F* (1, 68) = 7.75, *p* = 0.007, η_p_ ^2^ = 0.102] and a trial bin x previous trial structure interaction [*F* (7, 476) = 3.45, *p* = 0.001, η_p_ ^2^ = 0.048]. Again, the main effect of the previous rule was non-significant [*F* (1, 68) = 0.622, *p* > 0.250, η_p_ ^2^ = 0.009] as was the three-way interaction between trial bin, previous trial structure, and previous rule [*F* (7, 476) = 0.89, *p* > 0.250, η_p_ ^2^ = 0.013].

We also specifically compared positive transfer effects in the same rule and different rule groups. Log-transformed RT data from both training and test blocks of the CF -> CF (same rule), and CF -> CF (different rule) groups were analysed with a mixed-model with block (2 levels: training or test) and trial bin (8 levels: 1-8) as within-subject factors and previous rule (2 levels: same rule, different rule) as a between-subject factor. The ANOVA revealed significant main effects of block [*F* (1, 238) = 10.07, *p* = 0.003, η_p_ ^2^ = 0.229] and trial bin [*F* (7, 238) = 3.03, *p* = 0.005, η_p_ ^2^ = 0.082], and a significant block x trial bin interaction [*F* (7, 238) = 3.20, *p* = 0.003, η_p_ ^2^ = 0.086]. Importantly, the main effect of the previous rule was non-significant [*F* (1, 34) = 0.31, *p* > 0.250, η_p_ ^2^ = 0.009] while the three-way interaction between trial bin, previous trial structure, and previous rule was marginally non-significant [*F* (1, 68) = 1.99, *p* = 0.068, η_p_ ^2^ = 0.055]. Overall, this suggests that while prior rule experience may have some influence on subsequent performance, the positive transfer effect is largely driven by prior experience with a trial structure.

## Discussion

Psychologists have focused almost exclusively on the relations between stimuli, contexts, responses and outcomes as a framework for understanding cognitive control and our ability to adapt and generalize to novel task environments. In this paper, we developed the hypothesis that internal control policies, required for coordinating cognitive processing during a task, form an essential component of task knowledge independently of the stimulus-response (S-R) rule structure of the task. Thus, we sought to test whether control policies can be learned and transferred to novel task situations separate from S-R rules. We tested this hypothesis in the context of a working memory control task, leveraging previous findings that subjects learn different selective gating policies for context first or context last trial structures (Chatham et al., 2014). We argue that the results presented in this paper are clear evidence in support of the hypothesis.

First, the experiments provide evidence of negative transfer. Participants with prior experience with a CL block were slower to respond on the initial trials of a subsequent CF block compared to participants with no prior experience. This effect was not driven simply by WM load experienced during the CL block, as participants in the CF2 -> CF group, who were matched for load, did not show such slowing. Importantly, the negative transfer effects were asymmetric, in that they were not obtained in participants carrying out a CL block after having previously experienced a CF block. Thus, simply encountering a novel trial structure or a task switch was not a cause of the initial slowing. Rather, such an asymmetric effect is consistent with gating policy transfer, as the selective output gating policy learnt in a CL block can support performance on CF trials, but a selective input gating policy learnt in a CF block cannot support performance on CL trials. Finally, such transfer appeared to be deliberate, as participants with prior experience of both CF and CL trial structures were not slowed on the initial trials of a CF block that immediately followed a CL block. In other words, rather than simply perseverating with the last used policy, participants transferred the appropriate gating policy to the new task context.

Second, the pattern of target position effects we observed provided specific evidence of the transfer of a gating policy. As previously discussed, target position benefits are contingent on utilizing a selective input-gating strategy. Participants who had previously experience the CF trial structure showed a target position effect on the initial trials of a subsequent CF block, but not those who had previously experienced only the CL trial structure. Therefore, the early trial behaviour in the test block was consistent with the use of a gating policy tuned to the previously experienced trial structure. In other words, initial slowing observed in the test block of the CL -> CF group was due to negative transfer of a less efficient gating policy. Toward the end of the block, however, as they gained experience and the negative transfer effects on RT diminished, a robust target position benefit emerged. This result directly supports our contention that participants transferred a selective output-gating policy from the CL to CF block.

Our experiments also provided evidence for performance improvements that may reflect the positive transfer of a gating policy. In Experiment 1, participants in CF -> CF group were significantly faster in the first trial bin, and marginally more accurate overall in the test block as compared to the training block. Given that the S-R mappings were held constant, such an improvement may simply reflect practice with the S-R mappings. However, the interruption of a task by an intervening cue like an instruction screen is known to produce restart costs (Allport & Wylie, 2000; Poljac, Koch, & Bekkering, 2009). Such a restart cost is observed even when no other competing task has been instructed, and is considerably larger when other interfering tasks have been instructed (Allport & Wylie, 2000). Indeed, the restart cost can be equivalent in magnitude as the first trial RT when the task is first encountered (Allport & Wylie, 2000). Nevertheless, here we found that initial response times in the test block were faster than initial response times in the training block and no different from terminal response times in the training block. Therefore, it is also possible that the prior experience with a consistent CF trial structure in the training block allowed participants to learn a selective input gating policy which was transferred to the test block.

A key feature of Experiment 1 is that these learning and transfer effects occurred without any change in the prevailing S-R rule structure. To further test the independence of internal control policies from S-R rules, Experiment 2 independently manipulated prior experience with an S-R rule versus the trial structure. We found that both negative and positive transfer effects are primarily driven by prior experience with a trial structure, regardless of whether the specific rules were repeated or were new. Strikingly, participants were more accurate on the second CF block in the CF -> CF condition, even when they were required to use different (instructed) S-R rules in each block, thus providing stronger evidence that this improvement was due to the positive transfer of a gating policy. While more complex practice accounts cannot be ruled by this data, collectively these results are consistent with the transfer of a selective input-gating gating policy contributing to these improvements.

The results from Experiment 2 suggest that S-R rules and gating policies may be dissociable aspects of task knowledge and that gating policies are abstract entities, not tied to specific S-R mappings. However, we note that our rule change manipulation was minimal – only individual contextual and lower-level items in the task were changed, while keeping category structure intact. Therefore, generalization may alternatively reflect category knowledge rather than an abstract gating policy. It is plausible that a stronger manipulation of rule or overall task context would produce larger effects of prior rule experience that may interact with gating policy transfer. Indeed, previous studies have demonstrated transfer of task rules (Badre et al., 2010; Cole et al., 2011; Collins & Frank, 2013; Shanks & Darby, 1998). Future work will be necessary to define the precise relationship between the constructs of rules and gating policies.

Abstract gating policies that are separable from the content (like S-R mappings) have been invoked by two very different theoretical proposals to account for such generalization phenomena (Kriete, Noelle, Cohen, & O'Reilly, 2013; Taatgen, 2013) Kriete et al. (2013) developed a corticostriatal neural network model incorporating an indirection (‘pointer’) mechanism (implemented by a gating policy). In their model, such a mechanism supports the separate WM representation of ‘roles’ and their ‘fillers’. With such an architecture, it would be possible to learn and transfer policies independent of the specific ‘fillers’ though this has not been implemented in the context of WM gating. On the other hand, working within a symbolic, production system framework, Taatgen (2013) developed the primitive element theory wherein production rules are decomposed into simpler elements, some of which carry out the task-general job of routing information. Once such task-general rule elements are learnt, they can be used in other tasks independent of the task-specific elements. Such task-general rule elements responsible for routing information bear a close resemblance with abstract gating policies that we consider here. In addition, we suggest that generalization may also be achieved by modifying domain general properties of control like action-selection decision thresholds, WM input-gating thresholds, or arousal levels.

More generally, our results show that internal control policies carry an important form of task knowledge about the dynamics (order and timings of events) of the task, separate from the specific S-R mappings. Such dynamics dictate the order and timing in which different cognitive processes must be engaged. Control policies tuned to such dynamic structure can be transferred across task contexts. Our results add to a growing literature on *structure learning,* extending its principles to the domain of cognitive control policies (Braun, Mehring, & Wolpert, 2010; Collins & Frank, 2013; Gershman, Blei, & Niv, 2010; Huys et al., 2015; Rougier, Noelle, Braver, Cohen, & O'Reilly, 2005). Structure learning refers to our ability to identify and leverage invariant structure in the space of natural tasks to improve learning of novel tasks (Botvinick et al., 2009; Botvinick et al., 2015; Gershman & Niv, 2010; Tenenbaum, Kemp, Griffiths, & Goodman, 2011).

For example, Collins and Frank (2013) showed that human participants learn to represent ‘task-sets’ (sets of S-R mappings) as abstract entities that can be generalized (or re-used) across different contexts. In the domain of motor skill learning, Braun, Aertsen, Wolpert, and Mehring (2009) showed that humans are able to extract the invariant structure in the space of mappings between sensory inputs and motor programs for reaching behavior, thus improving performance on previously unseen motor tasks. Our results extend these ideas to cognitive control policies, demonstrating that humans exploit a different form of shared structure present in the task dynamics. A tendency to exploit such structure affords the opportunity for re-using previously learned control policies in novel task settings, thus reducing the time and effort required to adapt. On the flip side, such a tendency would also lead to poorer initial performance when the transferred policy is not a good fit for the dynamic structure of the new task.

Within the domain of working memory, the hypothesis that task regularities can be exploited for a more efficient use of a limited WM capacity has a long tradition beginning with the notion of ‘chunking’ (Miller, 1956; Simon, 1974) More recently, several researchers have demonstrated that humans learn to exploit statistical regularities in the space of to-be-remembered items to form compressed WM representations (Brady, Konkle, & Alvarez, 2009; Mathy & Feldman, 2012; see Orhan, Sims, Jacobs, & Knill, 2014 for a review), allowing an increase in the number of items than can be remembered. Our contribution to this literature is to demonstrate that humans also exploit regularities in the dynamics of the task (i.e. in the relative sequential position of the contextual cues and lower-level items in our task) and learn efficient, generalizable policies for controlling *access* to WM (selective input or output gating policies).

In this paper, we have chosen to interpret our results within the gating framework. However, our primary claim is that control policies carry structural knowledge about task dynamics. This claim is not tied to the gating framework exclusively. It is possible to interpret these results within other frameworks of WM or cognitive control. For instance, our notions of selective input v/s output gating policies bear some resemblance to the distinctions between proactive v/s reactive modes of cognitive control postulated within the dual mechanisms of control (DMC) framework (Braver, 2012). On this view, selective input-gating policies rely on a proactive mode in which a goal-relevant contextual representation is activated and sustained in the anticipation of future control demand. On the other hand, selective output-gating policies rely on a reactive mode in which the goal-relevant contextual representation is transiently activated when the control demand arises. From this perspective, our findings could also be interpreted as evidence for the transfer of a control mode across different task contexts. Note, however, that in the DMC framework, the transient and reactive engagement of control is triggered, ‘on-the-fly’, when current control demands increase. Our finding of transfer of a selective output-gating policy from the CL to CF task in our experiments strains against the notion of reactive control. Instead, our results are more consistent with the learning and transfer of a reactive control *policy,* regardless of the current control demands.

Similarly, our results may also be interpreted within the ‘attention to WM’ framework (Oberauer & Hein, 2012). On this view, WM is postulated to have a region of direct access that can represent multiple items or elements over a delay as bindings between items and retrieval cues. Input gating is equivalent to an attentional process that selectively encodes an item within the region of direct access by establishing such a binding. Output gating, on the other hand, is equivalent to shifting the ‘focus of attention’ to a single WM representation by priming the relevant retrieval cue. A gating policy, then, is equivalent to an attentional control policy. Within this framework, our results would imply that participants build control policies that are tuned to the dynamics of attentional allocation demands, and transfer them to other task contexts independent of the specific bindings between items and cues.

Several questions about the nature of gating policy transfer remain open. For instance, our results only demonstrate ‘near transfer’ of gating policies – i.e. over a short time scale and between very similar task contexts. The present results leave open whether humans transfer gating policies across very different tasks that require analogous selective use of working memory. Evidence of such transfer would lead to the hypothesis that people build repertoires of gating policies that can be re-used in different kinds of novel task contexts and store them over longer time scales. A related question concerns whether the transfer is flexibly and adaptively controlled. In previous work on the learning and transfer of abstract ‘task-sets’, for example, it has been argued that transfer is flexible in that participants make adaptive decisions about whether to re-use one of many previously learned task-sets or build a new one (Collins & Frank, 2013; Collins & Koechlin, 2012). Our finding that practice with the CF trial structure in an earlier block markedly attenuates the negative transfer effect on a later block hints that such transfer is deliberate, though more direct evidence would be required to conclusively demonstrate that such transfer is truly flexible.

In conclusion, our study highlights the importance of WM gating policies in particular, and control policies in general, as a key component of the task knowledge that supports flexible behavior and task generalization. Understanding the mechanisms supporting broad generalization of gating policies would be critical to explaining the remarkable flexibility of human cognition.

## Author contributions

A. Bhandari and D. Badre designed the study. A. Bhandari conducted the experiments and analyzed the data. A. Bhandari and D. Badre wrote the paper.

## Acknowledgements

We thank Ryan K. Fugate, Aja Evans, Celia Ford, and Adriane Spiro for assistance with data collection. This work was supported by grants from NINDS (NS065046) and NIMH (MH099078, MH111737) at the NIH, and a MURI award from the Office of Naval Research (N00014-16-2832).

## Appendix A

Verbatim instructions provided to participants for Experiment 1:

“The task you will be doing today involves responding to sequences of items being presented on the computer screen. Each sequence will consist of three items presented one after the other. Your job is to identify a target item from the sequence based on a couple of rules. (Show the rules)

These are the two rules. They are in the form of trees. Each tree has a number at the top and associated with each number are a pair of items. 11 goes with the letters A and G. 53 goes with the symbols, 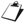 (dot) and π (pi). You need to memorize these relationships.

Now, the sequences that you will see will consist of 1 item from this pair (point to the letters pair), 1 item from this pair (point to the symbols pair), and 1 of the two numbers. For example, you might see the sequence A-Pi-11. Or the sequence might be 53-Dot-G. Your job is to identify the target item. The target item is always the item in the sequence that is associated with the number in the sequence.

(Show first example trial.) So, in the sequence A-π -11 – the target item would be A and not Pi, because A is associated with 11, while π is not. Similarly, in the sequence 53 – 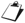 - G, the target would be 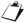. So, the number tells you which of the other two items is the target. Is that clear? (If not, repeat and clarify).

At the end of the sequence, just below the last item in the sequence, you will see a response panel. The response panel will have two items on the left and two items on the right. You have to look for the target item you have identified in this panel. If the target item that you have identified occurs as part of the left pair, you should press button 1. If the target item occurs as part of the right pair, you should press button 2. You should use the first two fingers of your right hand to make your response. You should make your response as quickly as possible without making errors.

The order in which the items are presented in the sequence is not important. Sometimes, the number might appear at the beginning of the sequence and sometimes it might appear at the end. Either way, the rules apply in exactly the same way. The target is always the item that goes with the number.

The items in the sequence each appear for a very short duration. The items will also be presented fairly quickly one after the other. So it is very important that you be alert and pay attention at all times. You will be doing a total of 2 blocks. It is important that you maintain a level of high alertness throughout the entire experiment.

Each block will begin with the rules being presented to you. You can review them for as long as you want. Once you are ready, press space bar to begin the block. There is no feedback, so make sure that you know the rules before you start the block. The rules are the same across all the blocks.

We will begin now. Remember to response as quickly as possible while being accurate.”

